# Microscale spatial fragmentation promotes bacterial survival under antibiotics

**DOI:** 10.64898/2026.03.17.712417

**Authors:** Dan Benbenisi, Tomer Orevi, Grayson S. Hamrick, Lingchong You, Nadav Kashtan

## Abstract

Most bacteria inhabit fragmented microscale environments rather than well-mixed systems in which antibiotic efficacy is typically defined. Here, we show that microscale spatial fragmentation alone can deterministically promote bacterial survival under antibiotic exposure. A coarse-grained theoretical model predicts that, at fixed bulk cell density and antibiotic concentration, fragmentation generates refuges through two coupled mechanisms: slower growth, which reduces susceptibility to growth-dependent antibiotics, and a finite antibiotic-per-cell constraint that lowers the effective intracellular antibiotic concentration. Using a microdroplet platform spanning five orders of magnitude in volume, we experimentally confirm these predictions for β-lactams; we further demonstrate that the same fragmentation-mediated protection extends to antibiotics with distinct modes of action. Spatial fragmentation thus creates robust, non-genetic antibiotic refuges governed by physical constraints rather than resistance or collective protection, with implications for bacterial persistence in natural and clinical environments and for antibiotic design.

## Introduction

Many microbial habitats are not continuous or well-mixed, but consist of innumerable, isolated or semi-isolated, microscale patches spanning several orders of magnitude in size. Such fragmented microscale habitats occur across diverse environments, including soil pores ^2–4^, leaf surfaces ^5–11^, and hydrated microdomains on skin ^12,13^ and in wounds ^14^, as well as tiny liquid residues on surfaces in the built environment ^15^. At these scales, spatial structure is not a secondary detail but a dominant physical constraint that directly shapes microbial population dynamics.

Microscale patchiness has been shown to strongly affect microbial interactions and competition outcome ^11,16^, population growth dynamics ^1^, community assembly and diversity ^17,18^, and horizontal gene transfer ^19^. Importantly, heterogeneity in patch size alone can generate large differences in local cell density ^1^, growth rate ^1^, and interaction strength ^16,18^, even in the absence of chemical gradients or genetic variation. Despite these advances, the ecological consequences of microscale fragmentation remain poorly quantified.

Bacteria are frequently exposed to antibiotics across both natural and human-associated environments. In soils and on plant roots and leaves, bacteria encounter diverse natural antibiotics produced by microorganisms and plants ^20–22^, as well as anthropogenic antibiotics introduced through agriculture and wastewater ^23–25^. Similarly, on human skin, in wounds, and on surfaces in hospitals and the built environment, bacteria are frequently exposed to topical antiseptics, residual antibiotics, and disinfectants ^26,27^. In all of these settings, antibiotic exposure occurs within thin hydration films, pores, or microscale liquid patches rather than in continuous, well-mixed volumes. Yet despite their pervasive exposure and strong selective effects, it remains unclear how microscale habitat structure, and specifically fragmentation and patch size distribution, modulates bacterial susceptibility to antibiotics across both ecological and clinical settings.

This raises a fundamental question: can microscale habitat fragmentation itself deterministically shape bacterial survival under antibiotic exposure, and how do patch size and isolation influence antibiotic efficacy?

Microfluidics and droplet-based studies have begun to reveal that compartment size itself can alter antibiotic action. Compartmentalization affects single-cell Minimal Inhibitory Concentrations (MIC) ^28,29^, and can alter antibiotic accumulation in cells ^30^. Work in droplet arrays has shown that antibiotic efficacy depends not only on nominal concentration but also on the number of antibiotic molecules available per cell, effectively linking killing to “antibiotic per cell” metric rather than bulk concentration alone ^31^. Under wet–dry cycles, transient microdroplet drying and wetting can protect bacteria from multiple antibiotic classes ^32^.

These observations echo the broader inoculum effect (IE), a long-recognized phenomenon in which higher initial bacterial density reduces apparent antibiotic efficacy ^33,34^. The inoculum effect is often attributed to collective protection such as aggregation or enzymatic degradation (e.g., β-lactamase activity). However, several studies indicate that purely physical or kinetic effects including limited antibiotic molecules per cell or altered intracellular antibiotic concentration, can also contribute ^30,35–37^. Even modest differences in cell density can markedly shift MIC estimates ^34^. Initial cell density can additionally influence growth productivity, thereby further modulating IE ^38^. Theoretical work has further suggested that in fragmented habitats, high density per unit volume, rather than physical crowding per se, may drive IE-like behavior, for example through accumulation of antibiotic-inactivating enzymes ^39^. Yet the field still lacks an integrated framework to determine whether microscale habitat fragmentation produces general and reproducible patterns of antibiotic susceptibility that are not specific to a particular antibiotic class or mode of action. This gap persists in large because most experimental systems lack direct, quantitative control over patch size while simultaneously generating patch-size distributions spanning several orders of magnitude within a single platform. Moreover, it remains unclear how multiple mechanisms, including density-driven molecule-to-cell ratios and confinement-induced growth slowdown, act together across a broad range of patch volumes.

Here we address this gap by combining theory with controlled microdroplet experiments. We developed a theoretical framework predicting that microscale confinement generates antibiotic refuges through two coupled mechanisms (Fig. 1A,B): (i) reduced bacterial growth rates in small droplets ^1^, decreasing susceptibility to growth-dependent antibiotics, and (ii) a density-driven effect arising from the finite ratio of antibiotic molecules to cells, which lowers target-site saturation even at fixed bulk concentration. The framework is grounded in the well-established growth-dependent action of β-lactam antibiotics ^40–44^. Both mechanisms emerge strictly from confinement, as even a few cells in a small droplet can reach high density per unit volume and experience diminished effective antibiotic load.

**Figure 1.**
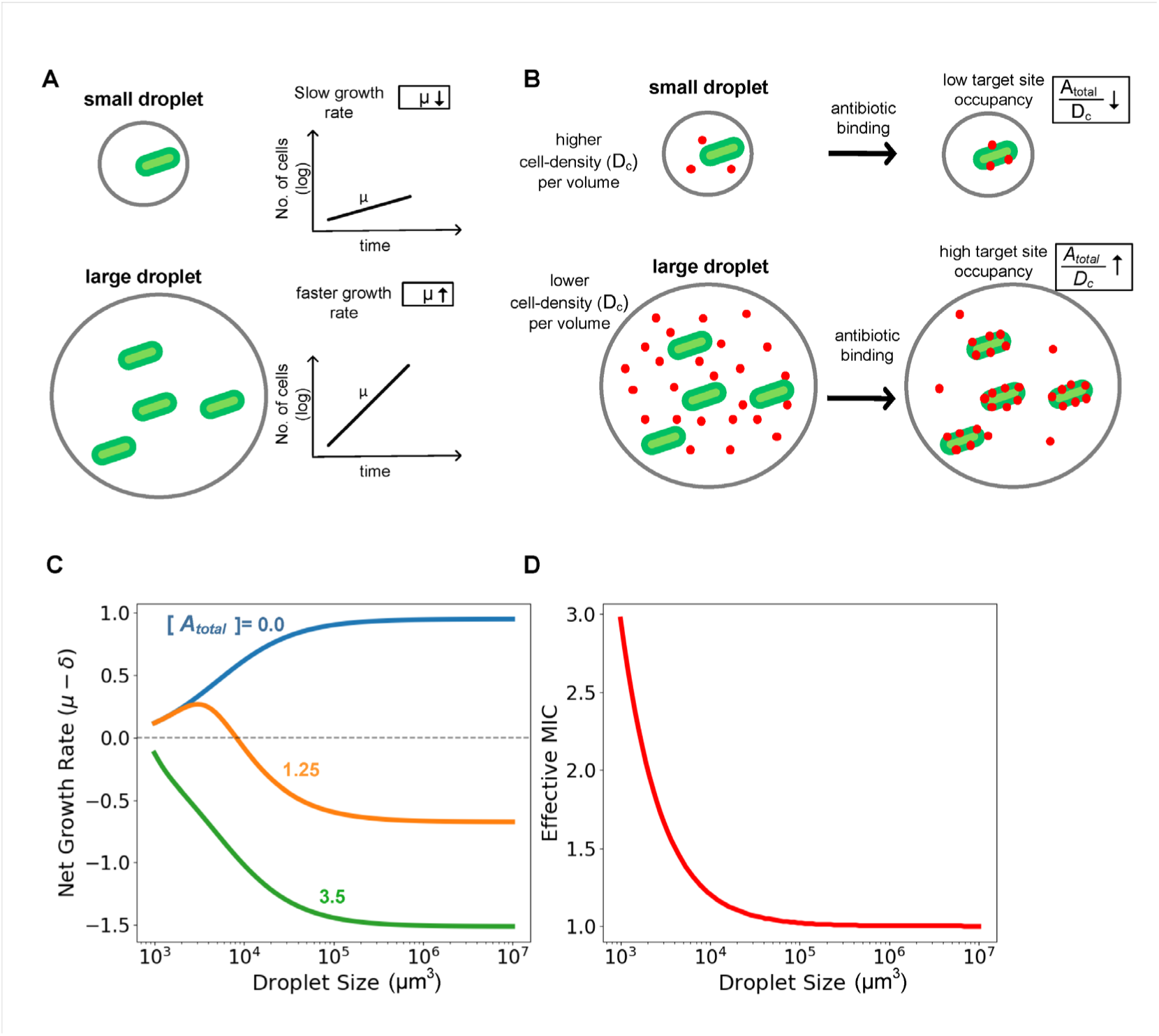
Theory predicts droplet-size-dependent survival under antibiotics. **(A)** Conceptual illustration of bacteria inhabiting droplets spanning several orders of magnitude in volume. Growth rates are lower in small droplets, as established by Mant et al. ^1^, thereby reducing susceptibility to growth-dependent antibiotics **(B)** At a fixed bulk antibiotic concentration, the number of antibiotic molecules available per cell is inversely related to cell density per unit volume, leading to lower target-site occupancy in small droplets. **(C)** Model prediction for the balance between growth and antibiotic-mediated killing (μ − δ) as a function of droplet volume. At intermediate antibiotic concentrations (≈ bulk MIC; orange line), cells in small droplets grow, whereas cells in large droplets transition from net growth to net killing. At high antibiotic concentrations (> bulk MIC; green line) the net growth is negative also for small droplets and further decreases with droplet size. **(D)** Predicted increase in effective MIC with decreasing droplet volume.

Using the µ-SPLASH platform ^1^, which enables the study of bacterial cells in droplets spanning several orders of magnitude in volume, we first experimentally test these predictions using the β-lactam ampicillin. We then examine whether the same qualitative behavior extends to antibiotics from other classes, including aminoglycosides and lipopeptides. Across all tested antibiotics, bacteria confined in small droplets survive antibiotic levels that are lethal in larger droplets. This effect arises in the absence of genetic resistance or collective antibiotic inactivation. Instead, it is an emergent consequence of microscale habitat fragmentation and local cell density.

## Results

### Theory predicts small droplets promote bacterial survival under antibiotics

Our platform generates droplets spanning a wide range of volumes (from 1 pL to 100 nL) and enables quantification of population dynamics in each droplet. We first focus on β-lactams, the most widely used antibiotics clinically and common in natural habitats.

In each droplet, population dynamics reflect the balance between growth and antibiotic-mediated killing (Fig. 1):

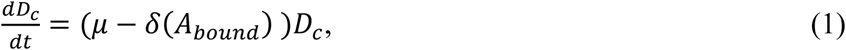

where 𝜇 is the effective growth rate, 𝛿 is the death rate, and 𝐴*_bound_* is the concentration of antibiotic bound to the target (e.g. penicillin-binding proteins for β-lactams ^45^, or ribosomes for aminoglycosides ^46^ or cell membrane for peptides ^35^).

In general, 𝛿 increases linearly with 𝜇 and nonlinearly with 𝐴*_bound_* ^47^ described by the following equation:

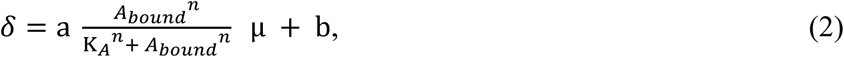

where K_𝐴_ is the threshold at which 𝛿 reaches half maximum, and *n* is the Hill coefficient (typically *n* >>1, producing a step-like threshold effect at K_𝐴_). The parameters *a* (typically >1) and *b* define the linear dependence of death rate on μ ^40^.

Droplet size modulates both μ and 𝐴_bound_. Smaller droplets operate closer to carrying capacity and thus exhibit reduced μ (Fig. 1A)^1^. At the same applied bulk concentration, high local cell density in small droplets reduces effective 𝐴_bound_ through titration (Fig. 1B; Methods).

Together, these effects predict a monotonic reduction of antibiotic efficacy with decreasing droplet size. This is captured by plotting the net balance μ − δ(𝐴_bound_) versus droplet volume (Fig. 1C). In the large-droplet limit, behavior approaches bulk culture: when bulk concentration slightly exceeds the MIC, killing dominates and μ − δ < 0. As droplet size decreases, reduced μ-dependence and lower 𝐴_bound_ progressively weaken killing, causing μ − δ to increase and eventually become positive below a critical droplet size. Thus, small droplets form survival regimes despite identical applied antibiotic concentrations. Increasing bulk antibiotic concentration shifts the curve downward but preserves the ordering across droplet sizes (Fig. 1C).

Equivalently, defining an effective MIC(*V*) as the bulk concentration satisfying μ(*V*) = δ(𝐴_bound_(𝑉)), the model predicts that effective MIC increases monotonically as droplet size decreases (Fig. 1D). Smaller compartments therefore require higher applied antibiotic concentrations to achieve the same lethal effect observed in large droplets.

These predictions depend only on coarse-grained quantities (μ and 𝐴_bound_) and do not rely on specific physiological mechanisms, establishing a simple and deterministic relationship between spatial fragmentation and antibiotic efficacy.

### Large-scale, high-resolution quantification of bacterial responses in fragmented microscale habitats

To test theory, we used the μ-SPLASH experimental platform (Fig. 2A)^1^. The system generates thousands of surface-confined microdroplets spanning five orders of magnitude in volume on a single “chip”, with fluorescently-tagged bacterial cells seeded stochastically across droplets. Time-lapse fluorescence microscopy enables tracking population dynamics in individual droplets (Fig. 2B, Fig. S1).

**Figure 2.**
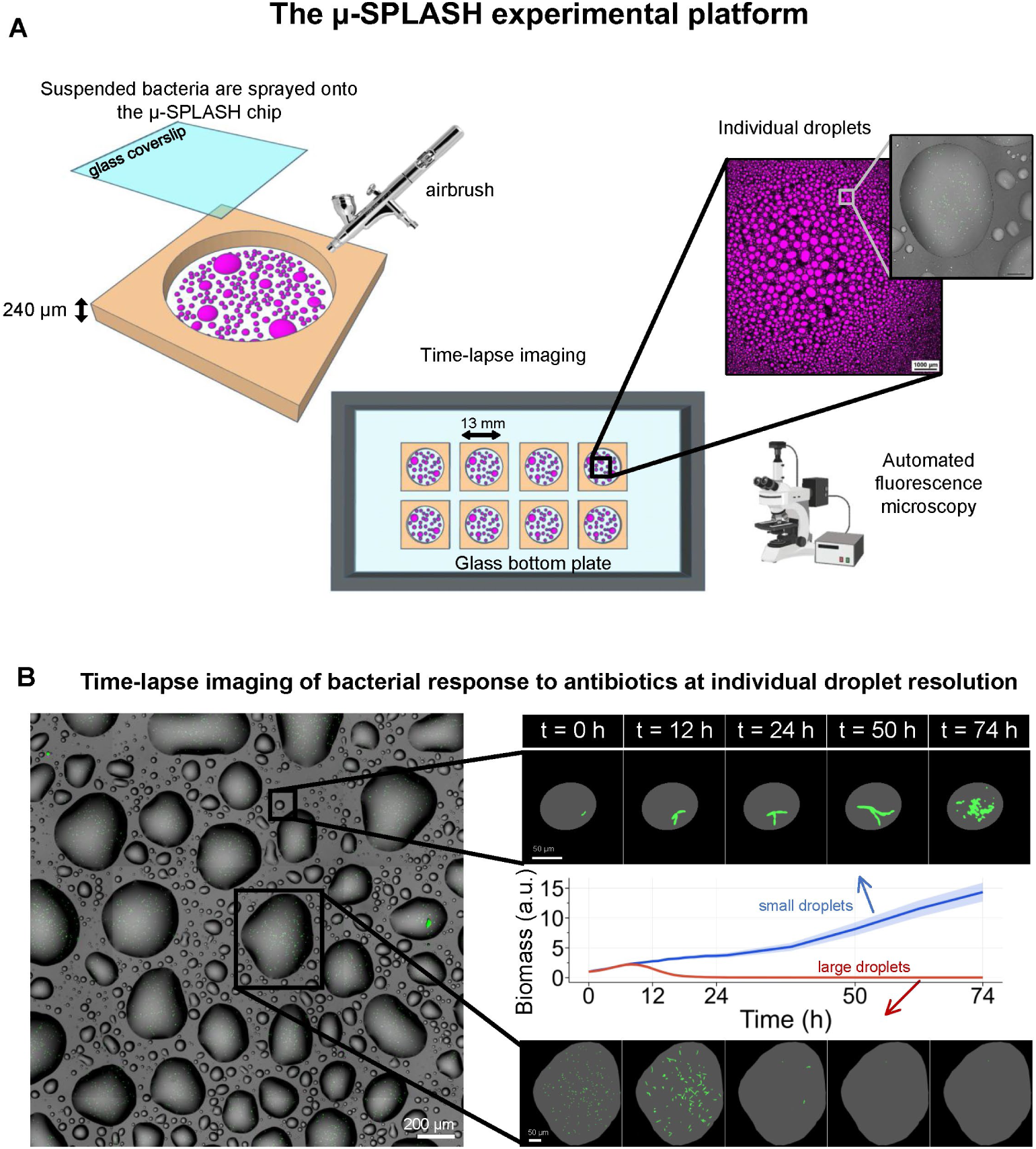
Large-scale quantification of bacterial responses in microscale fragmented habitats. **(A)** Fluorescently tagged bacteria are sprayed onto eight hollow adhesive spacers placed on a glass-bottom microplate using an airbrush. Each adhesive spacer (an individual chip) is sealed with a glass coverslip after spraying to maintain high humidity and keep the droplet intact. The plate is placed in a stage-top environmental chamber set at 28°C. Time-lapse imaging is conducted using a fluorescence microscope to track bacterial population dynamics. **(B)** Left panel: a representative section of a μ-SPLASH chip showing microdroplets spanning a wide size range (from 1 pL to 100 nL) and bacterial cells within (green dots). Right panel: Time-lapse microscopy sequences of *E. coli* responses to ampicillin at the bulk MIC in a small droplet (smallest size bin, 10³–10⁴ μm³) and a large droplet (>10⁶ μm³) over 74 h. The central graph shows the mean normalized biomass dynamics for these droplet-size ranges.

### Baseline population dynamics across droplet sizes

We first established baseline population dynamics of *E. coli* across droplet sizes in the absence of antibiotics. Bacterial biomass within each droplet was quantified as the area of the glass surface covered by cells (see Methods). For a direct comparison across droplets of different sizes, biomass dynamics in each droplet were normalized to their value at t = 0 h, highlighting the relative changes in biomass over time. Droplets were binned into four logarithmic volume ranges: 10³ -10⁴, 10⁴ -10⁵, 10⁵ -10⁶, and > 10⁶ μm³. For each bin, we calculated the mean biomass across all droplets at each time point, generating the average growth curves per bin (Fig. 3A). Consistent with previous work, smaller droplets showed lower normalized yield ^1^ (Fig. S1E). Fold-change analysis, defined as the log₂ ratio between the final and initial biomass, further highlighted the strong dependence of growth on droplet size (Fig. 3C).

**Figure 3.**
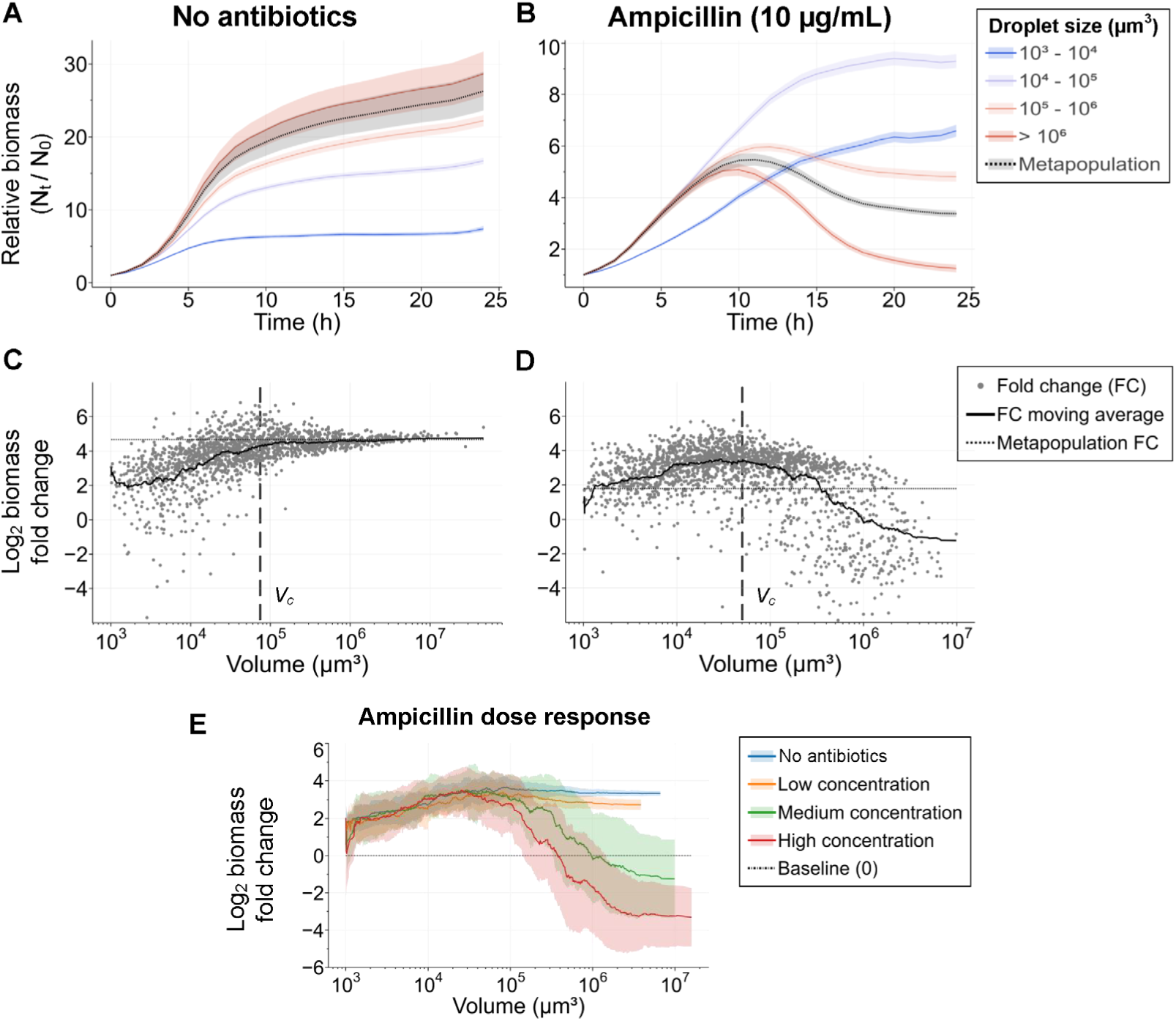
Small droplets promoted bacterial survival under ampicillin. **(A)** Mean normalized biomass dynamics across droplet size bins in the absence of antibiotics. **(B)** Biomass dynamics at the effective MIC of ampicillin (10 μg/mL). Small droplets (blue) maintain growth, whereas large droplets (red) show biomass loss. In large droplets, ampicillin triggers a short term response of biomass rise followed by a sharp decline, consistent with β-lactam–induced elongation followed by lysis. **(C,D)** Log₂ biomass fold change (final/initial) as a function of droplet volume without antibiotics (C) and under ampicillin (D). In the presence of ampicillin, fold change decreases sharply above the critical volume *Vc* (vertical line), indicating that cells in larger microdroplets are consistently more susceptible to antibiotic-induced biomass loss. **(E)** Graphs show the log_2_ biomass fold change (moving average ± SD) as a function of droplet volume. The blue line represents the antibiotic-free control, while the orange, green, and red lines represent concentrations below, at, and above the “effective MIC” (3.3, 10, and 30 μg/mL, respectively). Dose-response analysis across antibiotic concentrations shows progressively stronger inhibition with increasing antibiotic concentration and droplet volume as theory predicts (Fig. 1C,D).

### Testing bacterial response to antibiotics

Having the baseline droplet-size-dependent growth dynamics established, we analyzed the response of *E. coli* to ampicillin, a β-lactam that inhibits cell-wall peptidoglycan synthesis by binding penicillin-binding proteins ^45^. To quantify how antibiotic response depends on droplet size, we conducted μ-SPLASH experiments in which *E. coli* cells were exposed to ampicillin. Because MIC values measured in liquid cultures do not necessarily apply to microdroplets, we empirically determined an “effective MIC” by testing increasing concentrations until a clear inhibitory effect was observed in those largest droplets within a chip (> 10⁶ μm³). For Ampicillin we observed that the “effective MIC” was similar to the MIC in bulk culture (10 μg/mL).

### Droplet-size-dependent protection from ampicillin

We first examined bacterial responses at the empirically defined MIC of ampicillin (10 μg/mL). Biomass dynamics was analyzed using the same approach as in the antibiotic-free control, with droplets binned by volume, and mean biomass dynamics curves normalized to t = 0 h to assess relative biomass changes over time within each bin (Fig. 3B).

Consistent with model predictions (Figure 1C, D), small droplets showed increasing biomass over time, whereas large droplets exhibited growth arrest or biomass loss. In large droplets, ampicillin caused a rapid increase in biomass followed by a sharp decline, consistent with β-lactam-induced cell elongation followed by lysis ^42^ . In contrast, small droplets displayed modest but sustained biomass accumulation without detectable lysis (Fig. 3B), with a final yield only slightly lower than in antibiotic-free controls (Fig. 3A,C).

To assess longer-term dynamics, we extended imaging to 74 h (Fig. S2). Small droplets not only preserved biomass but frequently showed recovery and continued cell divisions, whereas large droplets displayed a persistent and continuous decline (Fig. 2B; Fig. S2A,C).

These droplet-size dependent patterns are also evident in the log_2_ ratio of final to initial biomass (Fig. 3D). We define a critical volume (*Vc*) as the droplet volume where the average cell density at t=0 h converges to the density of the source inoculum suspension ^1^. Droplets larger than *Vc* showed progressively lower fold changes, with negative values observed for volumes exceeding 10^5^-10⁶ μm³, indicating net biomass loss. Thus, susceptibility to ampicillin increased systematically with droplet size.

This droplet-size-dependence in bacterial response is antibiotic dose dependent (Fig. 3E, Fig. S3, S4). Consistent with model predictions (Figure 1C,D), the overall fold-change profile declines as antibiotic concentration increased. At the lowest concentration tested (3.3 μg/mL), growth was nearly indistinguishable from the antibiotic-free control. At the effective MIC (10 μg/mL) and above it (30 μg/mL), progressively stronger inhibition and biomass loss was observed.

### Mechanistic basis of ampicillin protection: growth rate and antibiotic binding

We next examined the mechanistic basis underlying droplet-size-dependent protection from ampicillin. With no antibiotics, instantaneous growth rates were consistently lower in small droplets than in large droplets (Fig. 4A), in agreement with our previous observation ^1^. Because ampicillin preferentially targets actively growing cells, reduced growth in small droplets is expected to lower lysis rates, providing a first mechanistic explanation for the observed protection in small droplets.

**Figure 4.**
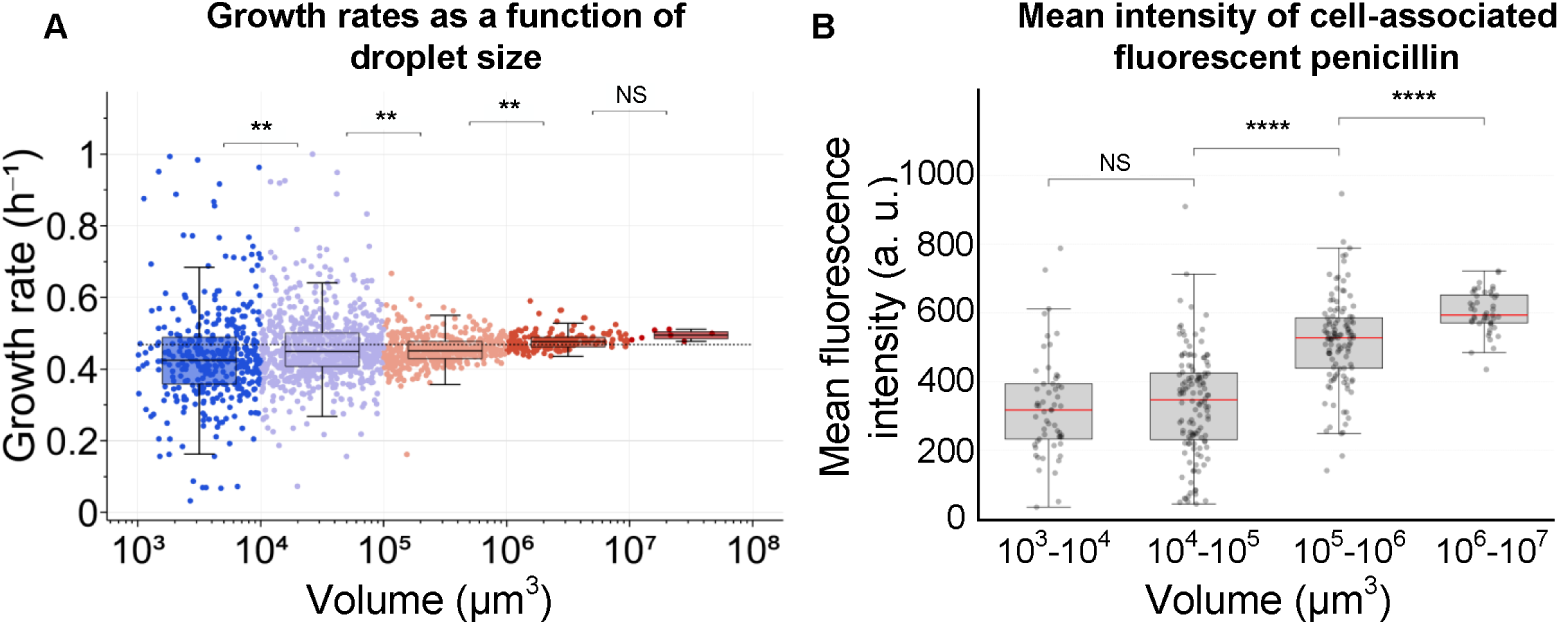
Mechanistic basis of droplet-size-dependent protection. **(A)** Mean growth rate as a function of droplet size. The mean growth rate without antibiotics is calculated based on the rate of biomass change in individual droplets over the first 4 h. **(B)** Fluorescent β-lactam binding assay (Bocillin FL) showing reduced cell-associated antibiotic in smaller droplets (see Methods). Each grey dot represents the mean intensity of a single droplet. Box plots show the median (red line) and the interquartile range. Small and intermediate droplets exhibit similarly low fluorescence, whereas large droplets (above *Vc*) show a sharp, statistically significant increase in fluorescence intensity, indicating higher target-bound antibiotic per unit biomass in larger droplets. Statistical comparisons in A and B are indicated above brackets (NS, not significant; **, p < 0.01;****, p < 0.0001).

In addition, cell density in the smallest droplets was approximately 20-fold higher than in the inoculum, and average density reached the inoculum level only in droplets with volume exceeding the critical volume (*V* > *Vc*) (Fig. S1C). Together, these observations support the two key components of our theoretical framework: slower growth and higher cell-density per unit volume in small droplets.

To directly test whether increased cell density in small droplets also reduces antibiotic availability at the cellular target level, we used a commercially available, fluorescently labeled penicillin (Bocillin FL ^48^), and quantified the mean intracellular fluorescence intensity as a measure of cell-associated, target-bound antibiotic across droplet sizes. Droplets in the small-to-intermediate size range had lower fluorescence intensities than droplets larger than *Vc* (≈10^5^ μm³), indicating lower levels of bound antibiotic per cell in smaller droplets (Fig. 4B, Fig. S5).

These results demonstrate that intracellular antibiotic levels are reduced in small droplets at the same applied bulk concentration, consistent with the model prediction that microscale confinement lowers target-site antibiotic occupancy. Taken together, the growth-rate measurements and Bocillin binding data provide direct empirical support that both reduced growth and reduced target occupancy contribute to protection against ampicillin in small droplets.

### Gentamicin and polymyxin B show similar droplet-size-dependent response

Having established droplet-size-dependent protection using ampicillin, we next asked whether the same protection extends to antibiotics with different cellular targets and growth dependencies. To test this we examined gentamicin, an aminoglycoside that binds the 30S ribosomal subunit and disrupts protein synthesis ^46^, and polymyxin B, a lipopeptide that disrupts Gram-negative membranes through interactions with lipopolysaccharide (LPS) ^49–51^.

The responses to gentamicin and polymyxin B at the “effective MIC” concentration revealed similar size-dependent patterns (Fig. 5A,B). Gentamicin induced a milder but consistent response, in which small and intermediate droplets exhibited growth though with somewhat lower yield relative to the antibiotic-free control, whereas the largest droplets showed growth arrest (Fig. 5A,C). Polymyxin B caused an immediate reduction in biomass in large droplets, while small droplets showed only weak inhibition and continued to accumulate biomass (Fig. 5B,D).

**Figure 5.**
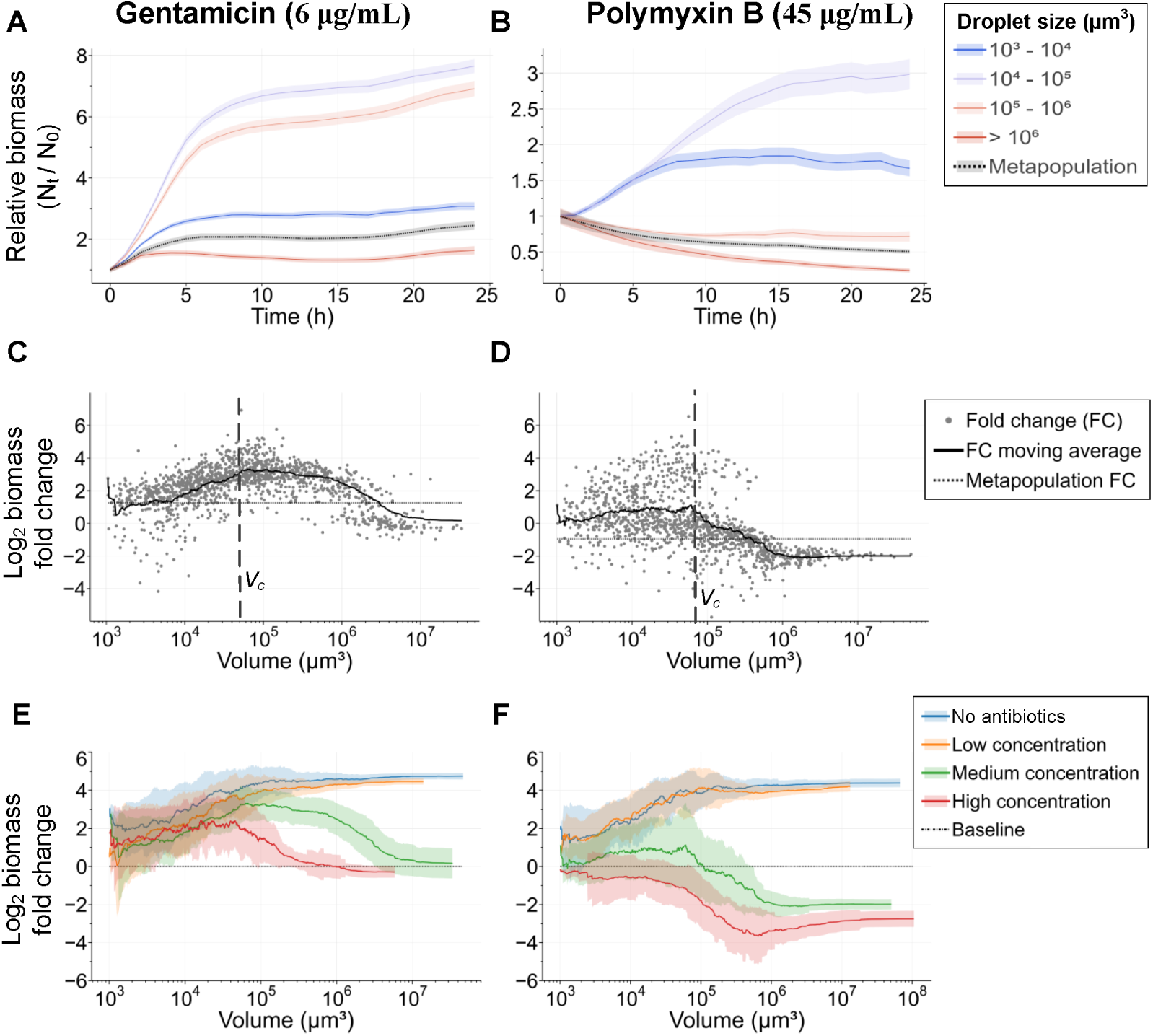
*E. coli* response to gentamicin and polymyxin B shows droplet-size-dependent protection similar to ampicillin. **(A,B)** Mean normalized biomass dynamics across droplet size bins at the effective MIC of gentamicin (A) and polymyxin B (B). Small droplets (blue) maintain biomass growth, whereas large droplets (red) show growth inhibition or biomass loss. **(C,D)** Log₂ biomass fold change (final/initial) as a function of droplet volume under gentamicin (C) and polymyxin B (D). Fold change decreases above the critical volume *Vc* (vertical line), indicating that cells in larger microdroplets are more susceptible to antibiotic-induced biomass loss. **(E,F)** Dose-response analysis across antibiotic concentrations showing progressively stronger inhibition with increasing antibiotic concentration and droplet volume. Graphs show log₂ biomass fold change (moving average ± SD) as a function of droplet volume for increasing concentrations of gentamicin (2, 6, and 18 μg/mL) (E) and polymyxin B (15, 45, and 135 μg/mL) (F). The blue line represents the antibiotic-free control, while the orange, green, and red lines represent concentrations below, at, and above the effective MIC, respectively.

Dose-dependent analyses of gentamicin and polymyxin B, based on experiments conducted below, at, and above the effective MIC, mirrored the trends observed for ampicillin (Fig. 5E, F). As antibiotic concentration increased, the overall fold-change profiles declined, indicating stronger inhibition. Importantly, this inhibitory effect became progressively more pronounced with increasing droplet volume, and the droplet volume at which biomass decline occurred shifted toward smaller volumes at higher antibiotic concentrations. Across all tested concentrations and antibiotics, cells in small droplets (*V* < *Vc*) were consistently more protected than those in large droplets.

To assess whether the density-driven mechanism also applies to polymyxin B, we performed experiments using fluorescently labeled polymyxin B ^52^ and quantified the mean cell-associated fluorescence as a proxy for cell-bound antibiotic across droplet sizes. As observed for the fluorescent β-lactam, average fluorescence per cell increased with droplet size (Fig. S6), indicating higher levels of membrane-bound antibiotic in larger droplets, in agreement with model predictions.

Unlike ampicillin and gentamicin, polymyxin B acts primarily by disrupting bacterial membranes rather than inhibiting growth-dependent biosynthetic processes, thus reduced growth rates are not expected to strongly decrease susceptibility ^51^. The persistence of strong droplet-size-dependent protection for polymyxin B thus suggests that the density-driven limitation alone is sufficient to protect cells in small droplets (see also Discussion).

### Antibiotic exposure reshapes metapopulation distribution, increasing the relative contribution of small droplets

Last, we examined how antibiotic exposure alters the population distribution across droplet sizes. We quantified the fraction of the total population contained within each droplet size bin at the start of the experiment (t = 0 h) and after 24 hours (t = 24 h) (Fig. 6A-D). This analysis revealed distinct patterns resulting from the differential response of cells to antibiotics as a function of droplet-size.

**Figure 6.**
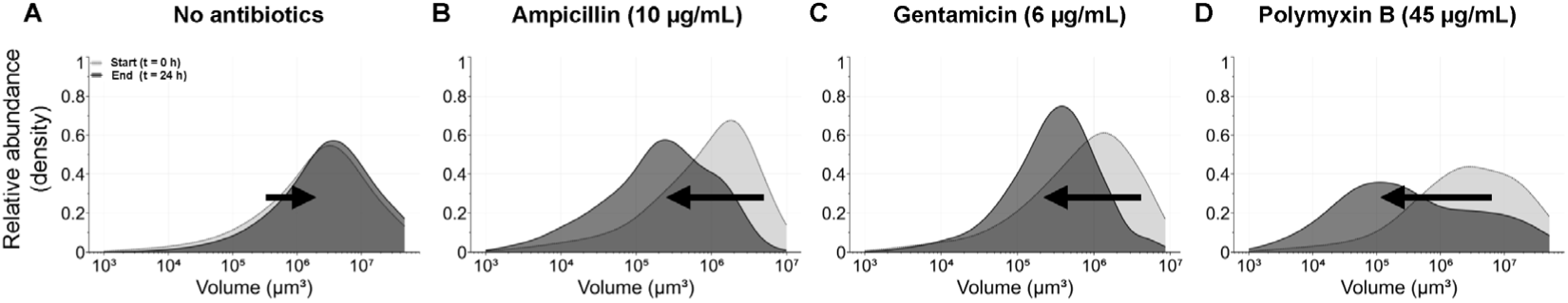
Antibiotic exposure redistributes bacterial metapopulation across droplet sizes. The fraction of the total bacterial population within each droplet size bin is shown at the start of the experiment (t = 0 h, grey shading) and after 24 hours (t = 24 h, black shading) for different treatments. **(A)** In the absence of antibiotic the population shows a modest shift toward larger droplets over time. **(B-D)** In contrast, antibiotic treatment exposure inverted this pattern: **(B)** ampicillin, **(C)** gentamicin, and **(D)** polymyxin B. All antibiotics caused a relative increase in the population fraction in smaller droplets and a decline in larger droplets.

Without antibiotics the metapopulation (the whole population within a single “chip”) experienced overall growth. This growth was accompanied by a small composition shift in the population, toward the large droplets ^1^ (Fig. 6A). In contrast, all antibiotic treatments resulted in a different pattern (Fig. 6B-D): the relative contribution of small droplets to the total population increased markedly, while the fraction of the population in large droplets declined significantly. This shift indicates that antibiotic exposure selectively reduces the population in large droplets, while sub-populations confined to smaller droplets increased in their relative abundance within the metapopulation.

## Discussion

Our theoretical framework predicted that microscale habitat fragmentation deterministically modulates antibiotic response in a patch-size-dependent manner, thereby generating antibiotic refuges. Two coupled mechanisms come into play: slower bacterial growth in small droplets and a reduced ratio of antibiotic molecules to cells resulting from high cell density per unit volume. Our experiments confirmed these predictions and revealed general, reproducible patterns that are not specific to a particular antibiotic or mode of action. Across three bactericidal classes, bacteria in small droplets repeatedly survived concentrations that caused biomass collapse in large droplets.

Quantifying growth dynamics in antibiotic-free controls verified the basis for the first mechanism, demonstrating that growth rates were consistently lower in small droplets. Using fluorescent β-lactam and fluorescent polymyxin B, we further provide evidence for the second mechanism, showing that cells in large droplets accumulated significantly higher levels of target-bound antibiotic per unit biomass than cells in small droplets. Notably, this physical “antibiotic-per-cell” limitation echoes the inoculum effect, yet here arises from micro-patch volume rather than physical crowding or collective protection mechanisms. Together, our results establish that microscale habitat structure alone can act as a deterministic physical filter on antibiotic efficacy, independent of genetic resistance ^53^, phenotypic heterogeneity ^54,55^, or enzymatic inactivation ^39,56^.

Our results demonstrate that droplet-size-dependent protection is a general phenomenon that holds across antibiotics with distinct modes of action, while also revealing antibiotic-specific modifiers of this response. For ampicillin and gentamicin, which preferentially target actively growing cells, both confinement-driven mechanisms are expected to act in concert: growth suppression in small droplets reduces susceptibility, and high cell density per unit volume lowers the effective antibiotic dose per cell. Polymyxin B, however, retains bactericidal activity against non-growing cells. The persistence of strong droplet-size dependence for polymyxin B therefore demonstrates that density-driven limitation alone is sufficient to produce pronounced protection in small droplets, independent of growth-rate effects.

It is important to note that stochasticity is evident in our data at the levels of both single cells and single droplets. Such variability is expected and likely reflects phenotypic heterogeneity, which is well documented, and is expected to have a stronger impact in small census populations such as microdroplets ^16^ and in single-cell responses to antibiotics ^54,55^. Consistent with this expectation, we observed substantial cell-to-cell variability in fluorescent antibiotic accumulation for both β-lactam and polymyxin B (Fig. S6,7). While this stochasticity contributes to droplet-level variability, it does not obscure the deterministic average patch-size-dependent trend observed across the metapopulation.

In addition to the two general mechanisms described above, polymyxin B is likely subject to an additional physicochemical effect that further amplifies droplet-size-dependent protection. Polymyxin B is an amphiphilic lipopeptide with surfactant-like properties and has been shown to strongly partition into lipid and hydrophobic interfaces ^52^. Based on these properties, polymyxin B is expected to accumulate at water–air interfaces within microdroplets, potentially reducing its effective concentration in the aqueous phase near surface-associated cells. Because bacteria in our system colonize the solid glass surface, such interfacial enrichment is expected to lower the local antibiotic dose they experience. This effect likely contributes to the substantially higher “effective MIC” of polymyxin B in microdroplets compared with bulk cultures. Moreover, because the surface-to-volume ratio increases steeply in small droplets, this interfacial sequestration should disproportionately protect cells in the smallest droplets, acting as an antibiotic-specific modifier that operates independently of, and in addition to, the two general confinement-driven mechanisms.

The microscale effects we report propagate beyond individual droplets. Antibiotic exposure consistently shifted the metapopulation toward small droplets, indicating selective elimination of cells in large patches and the persistence, and in some cases recovery, of subpopulations confined to small droplets. Thus, antibiotic exposure actively restructures fragmented populations rather than uniformly reducing population size. The macroscale outcome of antibiotic exposure therefore depends not only on molecular potency but also on the distribution of patch sizes, and in particular the frequency of small, isolated compartments, within a fragmented habitat.

Importantly, the protection observed here is fundamentally distinct from classical mechanisms of antibiotic tolerance and resistance. Unlike biofilm-associated protection (physical or physiological) ^57,58^, it does not require surface-attached multicellular structures or extracellular matrix production. Unlike aggregation-driven or enzymatic protection, it does not rely on collective antibiotic degradation or sequestration ^56,59^. And unlike genetic resistance, it arises on short timescales without genetic changes. Instead, protection emerges purely from physical habitat fragmentation and local cell density, creating refuges that persist under antibiotic exposure.

Overall, our results reveal a consistent droplet-size-dependent response across antibiotic classes, in which cells residing in small droplets display systematically higher antibiotic resilience than those in large droplets. Our theoretical framework, supported by experiments, suggests two general, coupled mechanisms that jointly underlie this antibiotic response. Microscale spatial fragmentation therefore generates robust, non-genetic refuges that preserve viable bacteria, and actively restructures populations toward small, confined compartments under antibiotic exposure. Recognizing and quantifying these microscale refuges is thus essential for understanding bacterial responses to antibiotics in fragmented habitats. Incorporating microscale habitat heterogeneity into ecological and clinical frameworks may therefore improve predictions of antibiotic efficacy, persistence, and surface decontamination outcomes. More broadly, our findings highlight microscale spatial heterogeneity as a critical but often overlooked axis in antibiotic action, and open new strategies to dismantle protective microhabitats and improve antibiotic performance across diverse environments, from environmental surfaces to host-associated habitats.

## Methods

### Theoretical model

#### Size-dependent growth rate 𝜇(𝑉)

We model bacterial growth as saturating with droplet size, reflecting increasing access to nutrients, oxygen, or space in larger compartments. The growth rate is given by

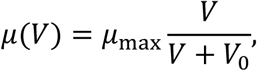

where 𝜇_max_ = 1 is the maximal growth rate in large droplets and 𝑉_0_ = 5,000 sets the characteristic volume at which growth reaches half-maximal levels. This functional form ensures that cells in very small droplets experience strong growth limitation, while growth asymptotically approaches bulk behavior at large volumes. The specific value of 𝑉_0_ was chosen to place the transition in growth rate within the experimentally relevant droplet size range, without introducing a sharp threshold.

#### Antibiotic sequestration and binding

Antibiotic availability is reduced by sequestration due to binding to cellular targets. For a fixed initial number of cells 𝑁, the effective cell density scales as

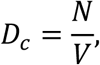

so that smaller droplets correspond to higher cell densities and therefore stronger antibiotic depletion.

For a given total antibiotic concentration 𝐴_total_, the free antibiotic concentration *A_free_*\* is determined implicitly by mass conservation:

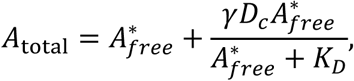

where 𝛾 = 0.045 is an effective sequestration strength per cell volume directly related to the concentration of target in the cell. 𝛾 can be considered a phenomenological parameter and was chosen such that antibiotic sequestration becomes appreciable only at high effective cell densities, while remaining negligible in large droplets, consistent with bulk measurements. Also, 𝐾_𝐷_ = 1.1 is the dissociation constant of high affinity antibiotic–target binding. This equation is solved numerically for *A_free_*\* at each droplet size.

The concentration of antibiotic bound to targets is then given by standard Michaelis–Menten–like kinetics,

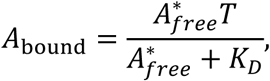

where 𝑇 represents the total target capacity, here normalized to 𝑇 = 1. Because 𝐷_𝑐_ increases as droplet size decreases, smaller droplets exhibit stronger antibiotic sequestration, leading to reduced free and bound antibiotic concentrations at fixed bulk dosing.

#### Pharmacodynamic killing

Cell death is modeled as a growth-coupled Hill-type response to bound antibiotic,

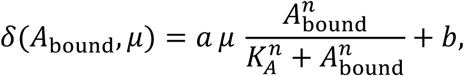

where 𝑛=10 is the Hill coefficient with a steep threshold, 𝐾_𝐴_ = 0.5 sets the sensitivity threshold, 𝑎 = 2.5 is the maximal antibiotic-induced killing rate relative to growth, and 𝑏 = 0.05 is a basal turnover rate. Coupling killing to 𝜇 reflects the assumption that actively growing cells are more susceptible to antibiotic action.

#### Net growth and effective MIC

The net growth rate is defined as 𝜇 − 𝛿. Figure 1C shows how net growth varies with droplet size for several fixed bulk antibiotic concentrations near the MIC. Figure 1D shows the effective MIC, defined as the minimal bulk antibiotic concentration required to drive net growth negative for a given droplet size.

Together, these simulations demonstrate that decreasing droplet size simultaneously suppresses growth and reduces effective antibiotic exposure via sequestration, leading to a strong and non-monotonic size dependence of survival and apparent resistance.

### Bacterial strains and growth conditions

*E. coli* MG-1655 cells harboring the pEB1-mGFPmut2 plasmid or the pEB2-mCherry2-L were used throughout the study (pEB1-mGFPmut2 and pEB2-mCherry2-L were gifts from Philippe Cluzel, Addgene plasmids #103980 and #104003 RRID:Addgene_103980_104003) ^60^. Culture media and growth conditions were as described previously ^1^ except that, in the final step, the optical density was adjusted to OD_600_=0.1. Antibiotics used in the study included: ampicillin (Bioprep, Israel), gentamicin (Bioprep, Israel), polymyxin B (Sigma), dansyl-labeled polymyxin B (Sigma) and BOCILLIN FL penicillin (Invitrogen, USA).

### µ-SPLASH experiments

A single-well glass-bottom plate (P01-1.5H, cellvis, USA) was used as a carrier vessel for eight µ-SPLASH chips. Each chip was constructed by stacking two adhesive imaging spacers (SecureSeal™ Imaging Spacer-SS1X13, grace biolabs, USA), creating a 13 mm-diameter, 0.24 mm-deep chamber (Fig. 2A). Excess margins of the 25 mm rectangular imaging spacer were trimmed to reduce chip dimensions to ∼20 mm, allowing eight chips to fit into a single well plate. Experiments were initiated by mixing *E. coli* MG1655 cells with the appropriate antibiotic to achieve cell density of OD_600_= 0.05 and the desired antibiotic concentration. Alexa Fluor 647 (Invitrogen) was added to a final concentration of 2 µM. Immediately after mixing, 100 µL of the suspension was transferred into the loading cup of an airbrush (IWATA model HP-M, Japan). The airbrush nozzle was pointed toward the center of a chip through a hollow cylinder (custom printed on an in-house 3D printer). The distance between the airbrush nozzle and the glass surface of the well was ∼5.5 cm. An air compressor connected to the airbrush was set to 1.5 bar and the fluid adjustment valve of the airbrush was tuned to level 4. A brief manual press on the airbrush activation lever was applied to deliver a sprayed sample onto the glass surface. Immediately after spraying, the chamber was sealed with a 18 x 18 mm glass coverslip (No.1, Marienfeld, DEU) on the upper, adhesive edge of the chamber. After all the chips were loaded the plate was mounted on a stage-top environmental chamber (H301-K-FRAME, Okolab srl, Italy) equipped with an open frame sample holder (MW-OIL, Okolab srl, Italy) and set to maintain 28 °C.

### Microscopy

Microscopic and image acquisition were performed using an Eclipse Ti-E inverted microscope (Nikon) equipped with Plan Fluor 10x/0.3 NA air objective. A LED light source (SOLA SE II, Lumencor) was used for fluorescence excitation. Dansyl polymyxin B fluorescence was excited with a 395/25 filter, and emission was collected with a T425lpxr dichroic mirror and a 460/50 filter. GFP or BOCILLIN FL fluorescence was excited with a 470/40 filter, and emission was collected with a T495lpxr dichroic mirror and a 525/50 filter. mCherry fluorescence was excited with a 545/25 filter, and emission was collected with a T565lpxr dichroic mirror and a 605/70 filter. Alexa 647 fluorescence was excited with a 620/60 filter, and emission was collected with a T660lpxr dichroic mirror and a 700/75 filter. All filters and dichroic mirror were purchased from Chroma, USA. A motorized encoded scanning stage (Märzhäuser Wetzlar, DE) was used to image multiple positions of the µ-SPLASH chips. The entire surface of the chip was scanned using the “large image mode” by imaging 10 x 10 adjacent fields of view (∼1.34 × 1.34 cm per scan). At t = 0 h (immediately after spraying), the droplet fluorescence signal was acquired by scanning the Alexa 647 channel for each of the eight chips. Next, bacterial fluorescence was imaged through the GFP channel by 11 z-planes (1.5 µm spacing; 15 µm total range) to ensure that all scanned sections of the chip are in the focus plane. Each chip was scanned hourly for 24 h followed by every 12 h for an additional 48 h. Images were acquired with an SCMOS camera (ZYLA 19 4.2PLUS, Andor, Oxford Instruments, UK). NIS Elements 5.02 (Nikon) software was used for acquisition.

### Dansyl polymyxin B and BOCILLIN FL penicillin binding assay

µ-SPLASH experiments were conducted in order to determine the binding level of two fluorescently labeled antibiotics to cells in microdroplets. Experiments followed the procedures described above, with the modifications detailed below. <u>BOCILLIN FL</u> <u>penicillin assay:</u> (a) overnight cultures were washed twice with 50 mM Tris buffer (Bioprep, Israel) and adjusted to OD_600_=0.5. To facilitate BOCILLIN FL entry across the cell envelope, cells were incubated with 2.5 mg/mL polymyxin B for 15 min, then washed twice with 50 mM Tris and filtered through a 5 µm disposable filter (Whatman, UK) to remove large cell aggregates. Immediately before spraying, cells were mixed with BOCILLIN FL and Alexa Fluor 647 to final concentrations of OD₆₀₀ = 0.15, 80 µM BOCILLIN FL, and 2 µM Alexa Fluor 647. (b) A Plan Apo 40x/0.9 NA air objective was used (c) imaging included 15 x 15 scans (covering 4.86 × 4.86 mm) capturing GFP and mCherry channels. Fluorescence signal was acquired through 7 z-planes (1.2 µm step; 7.2 µm range) to ensure in-focus imaging across all droplets. <u>Dansyl-polymyxin B assay:</u> (a) prior to spraying, bacteria were mixed with dansyl polymyxin B to achieve a final OD_600_=0.05 and 90 µg/mL dansyl polymyxin B. Alexa 647 was omitted in this assay. (b) a Plan Apo 20x/0.75 NA air objective was used (c) imaging consisted of 10 x 10 scans (covering 6.66 × 6.66 mm) capturing the bright-field, GFP, and dansyl polymyxin B fluorescence.

### Image processing and analysis

TIFF images were exported from the microscope software (NIS-Elements AR Analysis, version 5.02) in 16-bit RGB format. Image processing and analysis were performed using FIJI/ImageJ (version 1.54f) and Ilastik (version 1.3.3post3), together with customized Python scripts developed for data preprocessing and automated analysis.

For each chip, two fluorescence channels were extracted: GFP (25 time point images) and Alexa Fluor 647 (single time point). Bacteria segmentation was based on the GFP channel images and droplet segmentation was performed based on the Alexa channel images.

### Selection of best-focused z-planes from multi-z chip images

GFP fluorescence images were acquired as z-stacks of five focal planes. Because the optimal focal plane varied spatially across each chip, every image stack was tiled into 100 × 100-pixel sub-regions. For each sub-region, the corresponding five z-layers were evaluated using Laplacian-variance focus scoring to quantify image sharpness. The z-layer with the highest focus score was selected as the optimal plane for that sub-region. A composite best-focus image was then reconstructed by assembling all selected sub-regions into a single 2D representation.

### Droplet segmentation

For each chip, a droplet mask was generated using the Alexa Fluor 647 image acquired at the first time point. Segmentation was performed in FIJI/ImageJ using a custom macro. The workflow included contrast enhancement (CLAHE), Gaussian and median filtering, and conversion to 8-bit image followed by manual adjustment of the threshold to match the droplet contours. Holes within droplets were filled, and a watershed step was applied to split adjacent droplets. After manual inspection and correction, the resulting binary mask was saved as a TIFF file, and the droplet outlines were exported as ZIP file containing a distinct region of interest (ROI) for each segmented droplet. Additionally, a CSV file containing the droplets data (X,Y position, ID, and droplets pixel area) was exported. Because droplets remained largely static and stable throughout the experiment, this single droplet mask was applied to all subsequent time points. ROI overlays on the Alexa channel were used to visually validate segmentation accuracy

### Bacteria segmentation

The area of the glass surface covered by bacterial cells (derived from GFP signal) was used as a proxy for biomass. Bacterial biomass segmentation was performed using the Ilastik pixel-classification workflow, with the GFP fluorescence channel as input. For model training, a 7,000 × 7,000-pixel region was cropped from the center of each chip image and converted to HDF5 format. This provided a representative subset of image features while reducing computational load. Manual annotation of bacterial biomass vs. background was performed iteratively until the classifier reliably distinguished biomass from other image structures. Because phenotypic and physiological variations occurred across treatments, independent models were trained for each chip using nine representative images sampled at 3-hour intervals throughout the experiment. The trained model was then applied to the full-size GFP images (also converted to HDF5 format) to produce complete binary biomass segmentation maps. After training, all GFP images were batch processed, and the resulting binary masks were exported as TIFF files. For each time point, the masks were also converted into ZIP archives containing ROI files for each segmented biomass region. These ROIs were overlaid onto the original GFP images to visually confirm segmentation accuracy.

### Mapping bacterial biomass to droplets

After obtaining ROI files for both droplets and segmented bacterial biomass, bacterial biomass was mapped to individual droplets using a custom Python script. ROI metadata for droplets and biomass were extracted from the ZIP files and converted into polygon objects. For each time point, the script evaluated whether each bacterial ROI polygon was fully contained within a droplet polygon. If contained, the area of that bacterial ROI was attributed to the corresponding droplet. A minimum-area threshold was applied to exclude noise and very small artifacts from the analysis. The resulting dataset, containing the total bacterial area per droplet per time point, was exported as CSV files for downstream analysis.

### Droplet volume estimation

Python scripts were used to integrate bacterial biomass measurements with droplet-level metadata for each chip. Droplet properties were imported from CSV files and merged with biomass values per droplet obtained from the segmentation workflow. Spatial coordinates and areas were converted from pixels to micrometers using a scaling factor of 0.65 µm per pixel.

Droplet size, initially measured as projected area (µm²), was converted to volume by assuming a spherical-cap geometry. Droplet volume was estimated using the following equation ^61^:

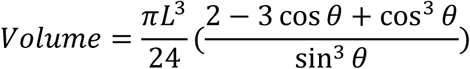

where L is the droplet diameter and θ is the contact angle. A previously measured contact angle of 32° (0.558 rad) for M9 medium droplets (∼350 µm diameter) on glass was used. Because the contact angle is not expected to vary substantially across the droplet size range studied (∼50–500 µm), this value was applied uniformly to all droplets.

### Bacterial biomass data processing and smoothing

Following volume estimation, bacterial biomass time series were preprocessed to correct artifacts and generate continuous growth profiles. Zero values at t = 0 h were replaced with NaN if later measurements indicated bacterial presence. Missing or zero values occurring between valid observations were interpolated using a geometric-mean approach, replacing each missing value with the geometric mean of the nearest preceding and following non-zero values. After automated processing, all time-series underwent a manual quality-control check to identify and correct segmentation artifacts, extreme outliers, or sharp discontinuities. Finally, bacterial biomass trajectories were smoothed using a LOWESS algorithm applied to log-transformed values, excluding the first and last two time points to minimize edge effects. The resulting smoothed dataset was exported for downstream analyses.

### Analysis of bacterial growth rates without antibiotics

Based on the biomass data, the slope of the change of log-transformed biomass over the first 4 h was calculated using linear regression, and used to determine growth rate in individual droplets (Fig. 4A).

### Visualization and inspection at individual droplet resolution

An interactive Python-based dashboard was developed to summarize the experimental data and enable real-time exploration of bacterial population dynamics across all chips and treatments. For each droplet, a time-lapse video was generated from the droplet ROIs and the corresponding fluorescence images, enabling direct visualization of growth (or death) patterns. These video clips were integrated into the dashboard, allowing simultaneous examination of quantitative metrics and visual growth outcomes.

### Statistical analysis

For selected statistical analysis, droplets were binned based on their volume on a logarithmic scale, using one-order-of-magnitude bin (10^3^–10^4^, 10^4^–10^5^, 10^5^–10^6^, 10^6^–10^7^, 10^7^–10^8^, µm³). Statistical differences between adjacent droplet-volume bins were assessed using permutation tests on the difference in means (or medians), with 1,000 random label permutations per comparison. Two-sided p values were computed from the empirical null distribution. Where multiple adjacent comparisons were performed, p values were adjusted using the Benjamini–Hochberg procedure.

### Quantification of cell-associated polymyxin B

Image analysis of the dansyl-labeled polymyxin B channel was performed to quantify the fluorescence intensity associated with bacterial cells within each droplet.

#### Focus selection

Each image was divided into 1,000 × 1,000-pixel sub-tiles, and the sharpest focal plane for each sub-tile was selected using the Laplacian-variance focus metric as described above. The resulting best-focus sub-tiles were stitched to reconstruct a single in-focus image.

#### Droplets segmentation

Droplet segmentation masks were generated in FIJI/ImageJ using the same workflow described in the Droplet segmentation section. Droplets missed by automated thresholding were manually added, by painting the missing regions, applying thresholding, and running Analyze Particles. The resulting binary mask and ROI set were used for all downstream analyses.

#### Bacterial segmentation

Bacterial regions were segmented using the Ilastik pixel-classification workflow following the same parameters described in the Bacterial segmentation section. To reduce noise, regions outside the droplet mask were excluded prior to classification, and segmentation was applied to the full image rather than cropped subsets.

#### Background correction and fluorescence extraction

Background fluorescence in the dansyl channel was corrected on a per-droplet basis using a Python-based polygon masking approach. For each droplet, the mean pixel value below the 75th percentile was calculated and subtracted from all pixel values within that droplet, reducing illumination and background variability. Within each segmented bacterial ROI, the mean of the top 25% brightest pixels was calculated and used as a proxy for labeled antibiotic within cells, per unit biomass. Qualitatively similar results were obtained when repeating the analysis with alternative parameter settings, including different intensity cutoffs and the use of median pixel values (for both background and intracellular fluorescence) instead of means.

The resulting intensity values were then aggregated at the droplet level to obtain the mean fluorescence per bacterium per droplet. Droplet-level intensity data were merged with droplet attributes, binned into logarithmic volume bins, and compared across bins. Statistical differences between neighboring bins were evaluated using two-sample t-tests with Bonferroni correction. Additionally, to assess the continuous relationship between antibiotic accumulation and droplet size, the Spearman’s rank correlation coefficient was calculated between the mean fluorescence intensity per bacterium and the log10 transformed droplet volume.

### Quantification of cell-associated Bocillin

Image analysis of the BOCILLIN FL fluorescence channel was performed to quantify antibiotic binding intensity using a workflow similar to the dansyl-polymyxin B analysis, but incorporating a refined local background-correction step. Bacterial biomass segmentation and droplet masks were obtained as described above. Droplet and bacterial segmentation were performed as described above. <u>Background correction</u> <u>and fluorescence extraction:</u> For local background estimation, a circular ROI with a radius of 50 pixels (∼8 µm) was defined around the centroid of each bacterial ROI. This ROI was intersected with the droplet mask to ensure that background estimates reflected only the local micro-environment within the droplet. The local background intensity was calculated as the mean of pixel values below the 75th percentile within this intersected region. This local background value was subtracted from the raw pixel intensities of the corresponding bacterium. Finally, to quantify the specific binding signal, the mean of the top 25% pixel-intensities within the background-corrected bacterial ROI was calculated. The resulting intensity values were aggregated per droplet and exported for downstream statistical analysis. Statistical differences between neighboring bins were assessed using two-sample t-tests with Bonferroni correction. Additionally, to assess the continuous relationship between antibiotic accumulation and droplet size, the Spearman’s rank correlation coefficient was calculated between the mean fluorescence intensity per bacterium and the log10 transformed droplet volume.

## Acknowledgements

AI tools (Gemini, ChatGPT) were used to assist with editing the manuscript for grammar and clarity. No AI tools were used for data analysis, figure generation, or interpretation of results.

## Author contributions

L.Y., T.O., and N.K. conceived the study. T.O. conducted experiments. D.B. performed image processing and data analyses. G.H., T.O., D.B., L.Y. and N.K. developed the theoretical framework. G.H. conducted mathematical modeling and simulations. All authors discussed the results and contributed to the final manuscript. L.Y. and N.K. supervised the project. D.B., T.O., L.Y. and N.K. wrote the manuscript.

## Data and materials availability

Data and Code are available at https://doi.org/10.6084/m9.figshare.31760281. All data needed to evaluate the conclusions in the paper are present in the paper and/or the Supplementary Information.

