## Supporting Information for "Microscale spatial fragmentation promotes bacterial survival under antibiotics"

**The supporting information includes Supplementary Figures S1-S6**

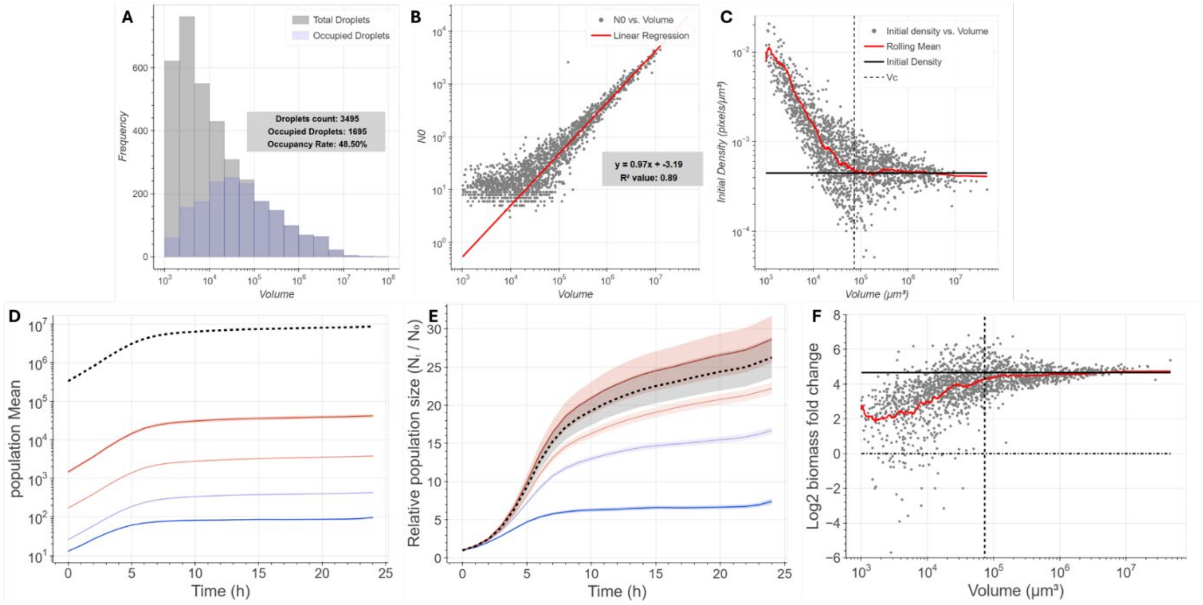


**Figure S1. *E. coli* population growth dynamics in the µ-SPLASH system in the absence of antibiotics.** Data are shown for a single representative control chip (no antibiotic) from the gentamicin experiment. (A) Histogram of droplet volumes, showing the distribution of all droplets (grey) and bacteria-occupied droplets (blue). Occupation probability increases with droplet volume. (B) Scatter plot of initial bacterial cell number ($N_0$) versus droplet volume at t = 0 h. The slope of the log–log relationship is ~1, indicating that cell seeding into droplets follows a Poisson process. (C) Initial bacterial density (cells per unit volume) as a function of droplet volume. Smaller droplets show higher densities, whereas larger droplets converge (red moving average) toward the bulk inoculum density at the critical volume (Vc, vertical dashed line). (D) Mean growth curves showing population size increase over 24 h for droplets grouped into four logarithmically spaced volume bins. Shaded regions indicate the standard error (SE). The dashed line denotes overall metapopulation growth. (E) Same as (D), shown with normalization to t = 0 h to highlight relative growth dynamics. (F) $Log_2$ fold change (final biomass relative to initial biomass) versus droplet volume over 24 h. Small droplets show lower mean fold change, which approaches the metapopulation value in droplets larger than Vc (red moving average).

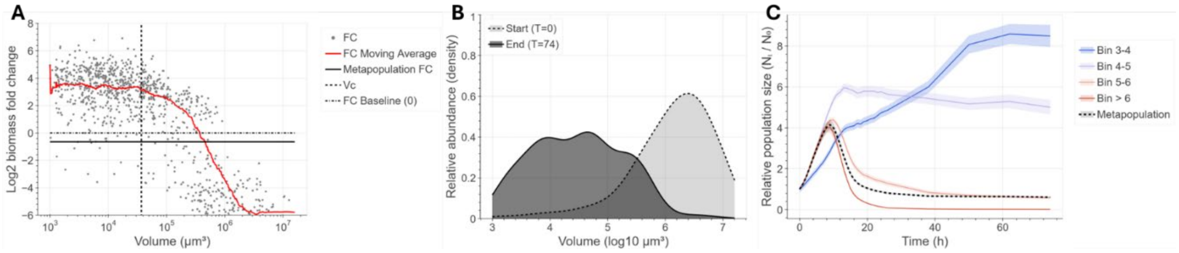


**Figure S2. Bacteria is small droplets show long-term resilience and viability following ampicillin treatment.** Data is based on imaging every 12 hours until 74 hours post-inoculation. (A) Fold-change analysis $Log_2$ of biomass at t = 74 h divided by biomass at t = 0 h. each dot represent an individual droplet. Large droplets show negative fold-change confirming their ongoing biomass collapse. Conversely, small droplets maintain consistent positive fold-change values, indicative of cell growth. (B) The final population distribution shows a drastic shift of the population toward small droplets. (C) Normalized growth curves show that populations in small droplets (V < Vc) remain viable and continue to accumulate biomass, while large droplets (V > Vc) display a continuous decrease. Shaded areas indicate standard error (SE).

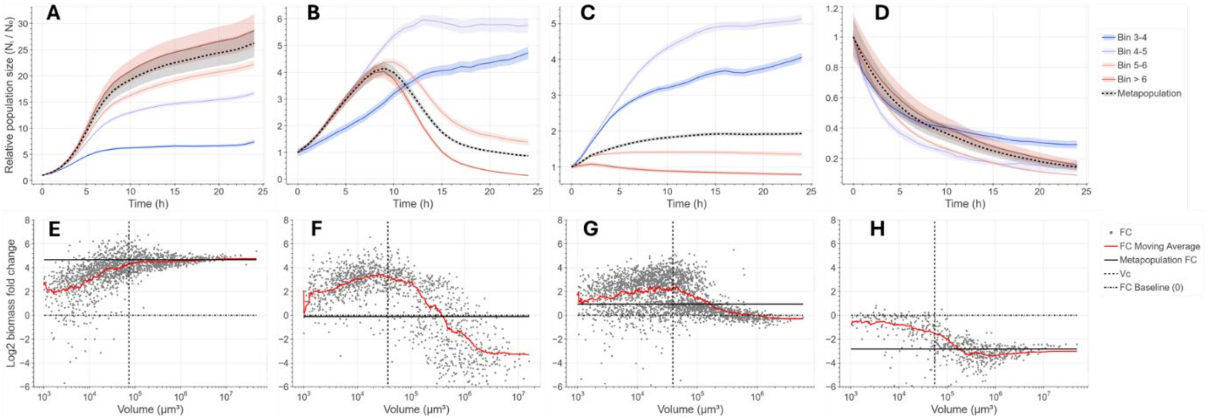


**Figure S3. Bacterial population dynamics at high antibiotic concentrations (above the effective MIC). (A–D)** Mean normalized biomass dynamics over 24 h for droplets grouped into logarithmic volume bins ($10^3$–$10^4$, $10^4$–$10^5$, $10^5$–$10^6$, $>10^6$ µm$^3$). **(A)** Control (no antibiotic). **(B)** Ampicillin (30 µg/mL), **(C)** Gentamicin (18 µg/mL), and **(D)** Polymyxin B (135 µg/mL). Compared to the effective MIC (Fig. 2), these higher concentrations induce stronger and more rapid biomass loss in large droplets (red lines). Despite this, droplet-size-dependent protection persists: small droplets (blue) maintain biomass or continue to grow, acting as antibiotic refuges even at these lethal antibiotic concentrations. **(E–H)** Biomass fold change (final/initial; FC) as a function of droplet volume. The volume above which biomass loss occurs is shifted toward smaller volumes compared to lower antibiotic concentrations (see Fig. 4). In (H) (Polymyxin B), the response is immediate and severe, with biomass loss spanning a broader range of droplet volumes than the observed at the effective MIC.

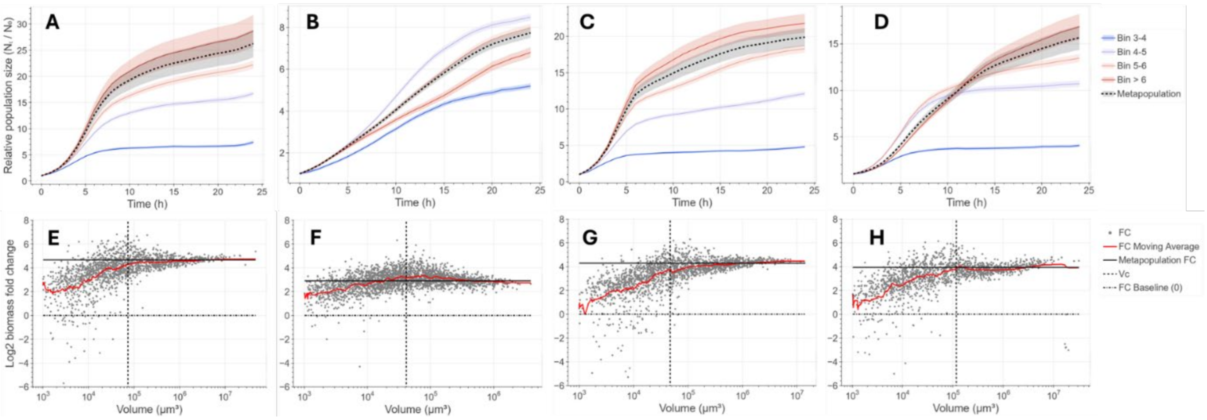


**Figure S4. Bacterial population dynamics under low antibiotic concentrations (below effective MIC). (A-D)** Mean normalized biomass dynamics over 24 h for droplets grouped into logarithmic volume bins ($10^3$–$10^4$, $10^4$–$10^5$, $10^5$–$10^6$, >$10^6$ µm$^3$). **(A)** Control (no antibiotic), **(B)** Ampicillin (3.3 µg/mL), **(C)** Gentamicin (2 µg/mL), and **(D)** Polymyxin B (15 µg/mL). In contrast to treatments at the effective MIC (Fig. 2), biomass dynamics across droplet sizes closely resemble the control (no antibiotics). In large droplets, biomass continues to increase but at a slightly reduced rate relative to the control. **(E–H)** Biomass fold change (final/initial; FC) as a function of droplet volume for the control and each antibiotic condition.

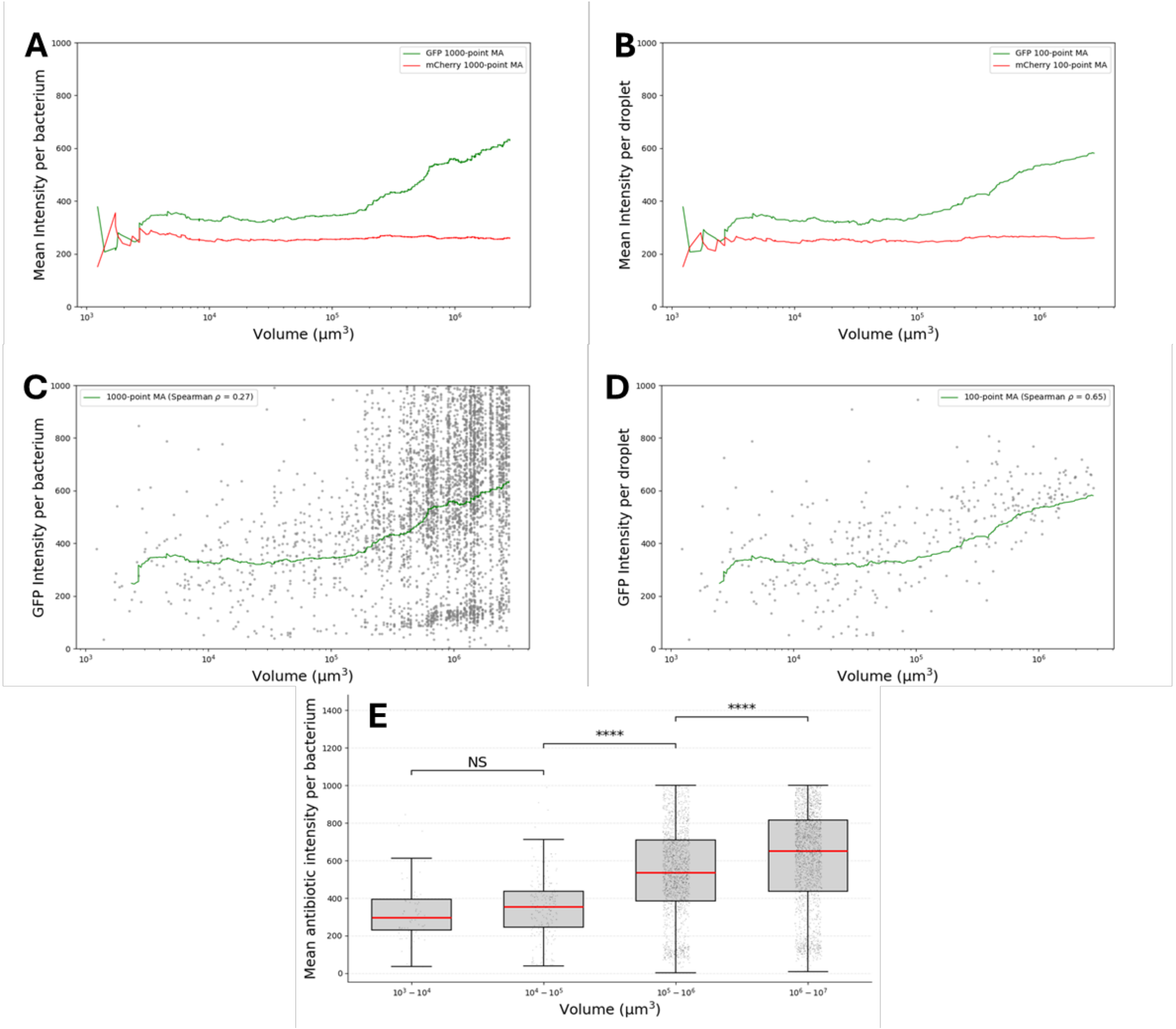


**Figure S5. Fluorescently-tagged penicillin shows higher cell-associated antibiotic levels in larger droplets. (A)** Moving average (1000 points) of fluorescence intensity per bacterium as a function of droplet volume for the antibiotic probe (Bocillin FL, green) and a constitutive marker (mCherry, red). **(B)** same analysis as in (A), averaged per droplet (moving average window is 100 droplets). The divergence between the two signals, where antibiotic binding increases with volume while the constitutive marker remains constant, indicates that the observed trend does not arise from optical artifacts or general physiological (e.g., cell size or protein density) but reflects specific antibiotic accumulation. **(C)** Moving averages of the Bocillin FL intensity as in (A) overlaid with single-bacterium data. **(D)** Moving averages of the Bocillin FL intensity as in (B) overlaid with individual droplet data. **(E)** Mean Bocillin FL intensity per bacterium (complementing the per-droplet analysis in Fig. 3B). Droplets were grouped into logarithmic volume bins, and statistical comparisons were performed as in Fig. 3 (NS, not significant; ****, $p < 0.0001$). The data confirm that bacteria in larger droplets ($10^5$ μm³ and above) show significantly higher cell-associated antibiotic signal (intracellular or target bound) than those in small droplets.

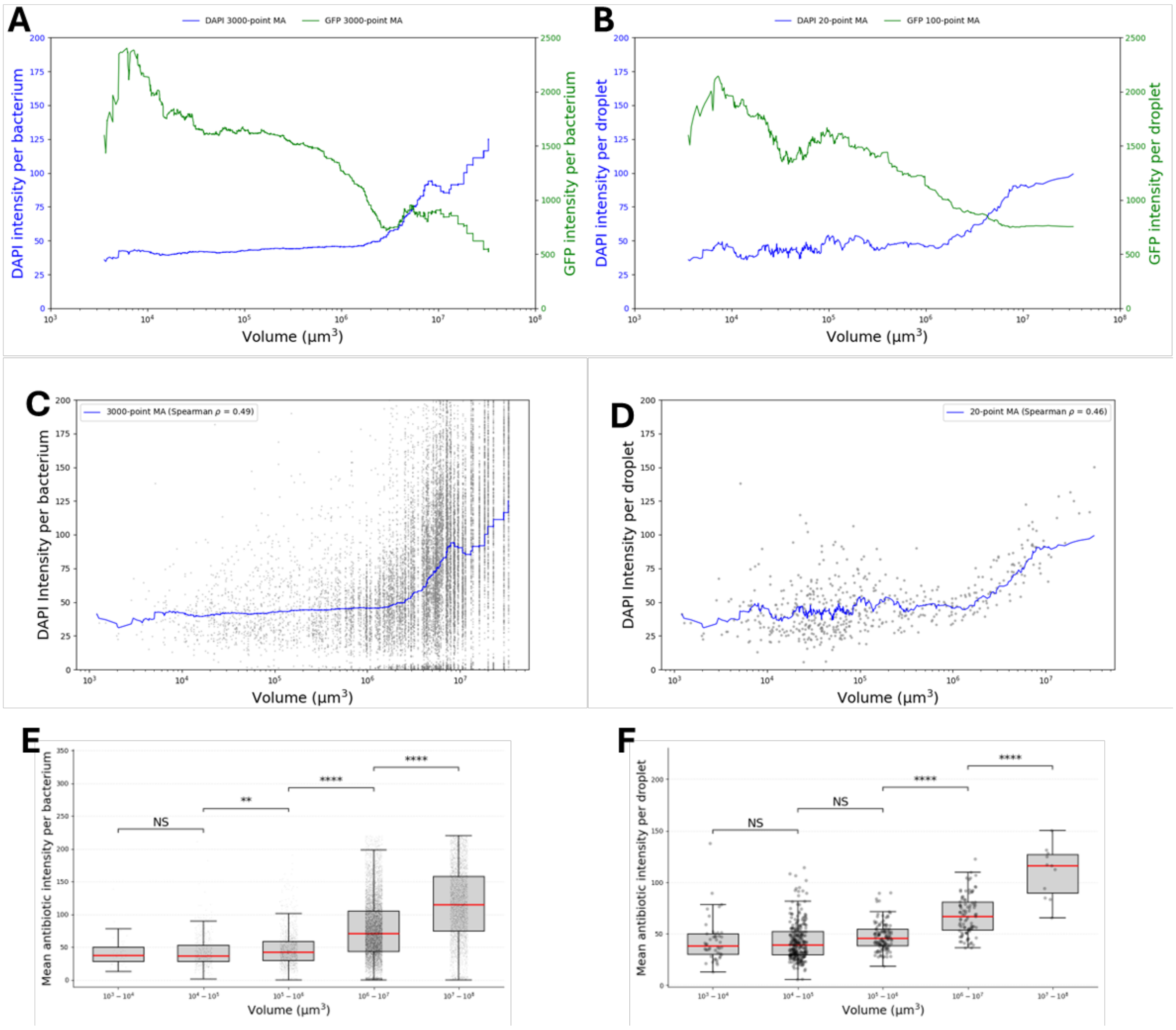


**Figure S6. Fluorescently tagged polymyxin B shows higher cell-associated antibiotic levels in larger droplet. (A)** Moving average (1000 points) of fluorescence intensity per bacterium as a function of droplet volume for the antibiotic probe (Dansyl–Polymyxin B, imaged in the DAPI channel, blue) and a constitutive marker (GFP, green). **(B)** Same analysis as in (A), averaged per droplet. (moving average window is 100 droplets) The divergence between the two signals, increasing Dansyl–Polymyxin B signal with droplet volume while the constitutive marker remains constant, indicates that the observed trend does not arise from optical artifacts or general physiological effects (for example cell size or protein density), but instead reflects specific antibiotic accumulation. **(C)** Moving averages of the Dansyl–Polymyxin B intensity as in (A) overlaid with single-bacterium data. **(D)** Moving averages of the Dansyl–Polymyxin B intensity as in (B) overlaid with individual droplet data. **(E)** Mean Dansyl–Polymyxin B intensity per bacterium. Droplets were grouped into logarithmic volume bins, and statistical comparisons were performed as in Fig. 3 (NS, not significant; ****, $p < 0.0001$). **(F)** Same as in (E) but data is averaged per droplet. Bacteria in larger droplets ($10^6$ μm³ and above) show significantly higher cell-associated antibiotic signal (membrane-bound) than those in small droplets.
